# Life history traits and a method for continuous mass rearing of the planthopper *Pentastiridius leporinus*, a vector of the causal agent of syndrome “basses richesses” in sugar beet

**DOI:** 10.1101/2022.05.07.491010

**Authors:** René Pfitzer, Mark Varrelmann, Klaus Schrameyer, Michael Rostás

**Affiliations:** Institute of Sugar Beet Research, Holtenser Landstraße 77, 37079 Göttingen, Germany; Victoriastraße 12/1, 74613 Öhringen, Germany; Agricultural Entomology, Department of Crop Sciences, University of Göttingen, Grisebachstrasse 6, 37077 Göttingen, Germany

**Keywords:** Cixiidae, *Candidatus* Arsenophonus phytopathogenicus, proteobacterium, development time, insect culture, mass rearing

## Abstract

**BACKGROUND:** The planthopper *Pentastiridius leporinus* (Hemiptera: Cixiidae) is the main vector of the γ-3 proteobacterium ‘*Candidatus* Arsenophonus phytopathogenicus’ which causes the syndrome “basses richesses” (SBR) in sugar beet. SBR is a new and fast spreading disease in Central Europe that leads to high yield losses. To date the development of management strategies is hampered by insufficient knowledge about general life history traits of the planthopper and, most importantly, the year round availability of insects reared under controlled conditions. Rearing of *P. leporinus* has been considered challenging and to date no protocol exists.

**RESULTS:** Here we describe a method for mass rearing *P. leporinus* on sugar beet from egg to adult, which has produced five generations and >20,000 individuals between June 2020 and March 2022. An alternative host such as wheat is not necessary for completing the life cycle. No-choice experiments showed that *P. leporinus* lays 139.1 ± 132.9 eggs on sugar beet, whereas no oviposition was observed on its nymphal host wheat. Head capsule width was identified as a trait that unequivocally distinguished the five nymphal instars. Developmental time from first instar to adult was 193.6 ± 35.8 days for males and 193.5 ± 59.2 days for females. Infection rates of adults were tested with nested polymerase chain reaction (PCR). The results demonstrated that 70-80% of reared planthoppers across all generations carried the SBR proteobacterium.

**CONCLUSION:** The mass rearing protocol and life history data will help overcome an important bottleneck in SBR research and enhance efforts in developing integrated pest management tools.

## 1 INTRODUCTION

The planthopper *Pentastiridius leporinus* (Linnaeus 1761) is a member of the Cixiidae (Hemiptera, Fulgoromorpha) family, which comprises almost 2,000 species worldwide.^1^ The insect has a palearctic distribution but does not occur in the northern regions.^2^ In natural ecosystems, adult *P. leporinus* are known to feed on aboveground parts of reed grass (*Phragmites australis*) but no information is available on host plants used by the immature stages that ingest phloem sap from roots.^1-3^ In agricultural systems, *P. leporinus* feeds on sugar beet (*Beta vulgaris*) leaves, thereby transmitting the phloem-restricted γ-3 proteobacterium ‘*Candidatus* Arsenophonus phytopathogenicus’ that causes a disease known as syndrome “basses richesses” (SBR).^4-6^ In recent years, the planthopper has become a major economic pest in several Central European sugar beet growing regions, including eastern France, Germany, and Switzerland.^7-9^ During the summer, adult *P. leporinus* migrate to sugar beet fields, where the females oviposit into the soil. The hatching nymphs start feeding on sugar beet roots but usually complete their development on winter wheat, which frequently follows in crop rotation. In the subsequent year, adult planthoppers emerge from the soil, climb up on wheat plants and then migrate into a nearby sugar beet field.^4,10,11^

Interestingly, *P. leporinus*, is not very common in natural ecosystems and in Germany it is even red-listed as an endangered species.^12^ A recent host plant shift to sugar beet and cereals such as winter wheat (*Triticum aestivum*) or barley (*Hordeum vulgare*) has probably led to the tremendous increase in population sizes of *P. leporinus* and thus to the fast spread of SBR.^4,10,11^ The sugar beet disease SBR leads to severe economic losses due to low sugar contents and yield reductions of up to 25%.^6,9,13^ Typical symptoms are chlorosis and necrosis of the older leaves, asymmetric young leaves and necrosis of the vascular bundles of the tap roots.^6^ SBR was first recorded in eastern France in 1991 and detected in Germany in 2009.^9,14^ More than a decade later, the area of affected sugar beet has accumulated to 3,000 ha (2019) in Switzerland and 16,400 ha (2018) in Germany.^7,8^

Apart from the γ-proteobacterium *Ca*. Arsenophonus phytopathogenicus, the phytoplasma ‘*Ca*. Phytoplasma solani’, belonging to the stolbur group (16SrXII), has also been identified as causal agent of SBR symptoms.^4,6,15-17^ However, so far only the proteobacterium plays a major etiological role in France and Germany.^6,8,9^ Adult *P. leporinus* and nymphs can transmit the SBR proteobacterium persistently to sugar beets.^18^ Furthermore, adult *P. leporinus* can also vertically transmit the proteobacterium to their offspring.^18^ Whether these microbes benefit the insect and its spread, is still an open question.

So far, no management strategies are available to farmers for controlling SBR and its insect vector. Due to the high mobility and the extended migration period of the adult planthopper, chemical control with available insecticides has not been practicable.^8^ While crop rotation and changes in soil tillage practices may show some promise and could be part of a future control strategy, further research is urgently needed.^10^

An important prerequisite for accelerated research on the biology, life history and management of *P. leporinus*, is a laboratory rearing that can provide a constant and sufficient supply of all developmental stages of the insect. Until now, continuous rearing of *P. leporinus* has been considered challenging and no protocol has been available.^18,19^ Here, we provide a method for the year round mass production of *P. leporinus* from egg to adult on sugar beet. In addition, data on various life history traits are presented. The information will enhance efforts in studying this economically important vector and its bacterial symbionts.

## 2 MATERIALS AND METHODS

### 2.1 Plants

Seeds of sugar beet, *Beta vulgaris* cv. Vasco (SESVanderHave Deutschland GmbH, Eisingen, Germany) without seed coating were grown in different pots and substrates, depending on the experiments. Plants were cultivated in a greenhouse with a temperature range of 20-35 °C. Natural daylight was supplemented with artificial light at 85 µmol (s m²)^-1^ (‘Elektrox SUPER BLOOM HPS 400 Watt’, Grow In AG, Berlin, Germany) when needed to maintain a 16:8 h light/dark photoperiod. A 3:1 mixture of peat (‘Fruhstorfer Erde Typ P 25’, HAWITA Gruppe GmbH, Vechta, Germany) and sand (0-2 mm diameter) was used as substrate. Plants were fertilized with a solution of 1 g l^-1^ ‘Hakaphos® Blau 15-10-15(+2)’ when necessary (COMPO EXPERT GmbH, Münster, Germany).

### 2.2 Insects

Adult *P. leporinus* were collected with a sweep net from a sugar beet field close to Neckarsulm in Germany (49°12’17.2”N 9°10’48.8”E) on 19 - 20 June 2020. Insects were identified according to their scutellum, vertex, pronotum, hind tarsus and male genital structures.^2^

If not stated otherwise, all experiments were carried out in a controlled environment room at 20.9 ± 1 °C, 48 ± 12.2% r.h. and a photoperiod of 16:8 h light/dark. Light intensity was 80 µmol (s m²)^-1^, provided by full-spectrum LED lights (‘Bioledex GoLeaf E2 LED Pflanzenleuchte Vollspektrum 120cm 50W IP44’, DEL-KO GmbH, Germany). Host plants and insects were cultivated in 60 × 60 × 60 cm cages (mesh size: 150 µm, ‘BugDorm-2120F Insect Rearing Tent’, MegaView Science Co., Ltd., Taiwan). Plants were only watered when showing very first signs of wilting to prevent planthoppers from drowning. Water was filled into the saucers, thus leaving the top part of the soil dry.

### 2.3 Role of plant container, substrate and environmental conditions for oviposition

As a first step in establishing a continuous rearing, the role of two different plant containers and three substrates for oviposition success was investigated in two environments. Transparent polystyrene cylinders (‘small pots’, 170 ml volume, 4.8 cm diameter, 10.4 cm height, ‘Zuchtgläschen’, K-TK e.K., Retzstadt, Germany) with a hole (1 cm diameter) in the bottom and 2.4 l polypropylene plant pots (‘large pots’, 16 cm top diameter, 15 cm height) were used. Both container types were filled with either field soil (loamy clay), a mix of peat and sand (3:1) or peat mixed with cracked expanded clay (‘Original LamstedtDan’, 4–8 mm, Fibo ExClay Deutschland GmbH, Lamstedt, Germany) (3:1). Field soil was collected from the sugar beet field where planthoppers were caught. The soil was heat-treated (45°C) for 48 h before use, in a ‘Kempson’ apparatus.^20^ A single sugar beet plant (small pots: growth stage according to BBCH 14, large pots: BBCH 16) was grown in each pot.^21^ Sugar beets in small pots received six female and three male adults, while plants in large pots received ten female and five male adults. A transparent, perforated polypropylene bag (small pots: 25 cm length × 15 cm width, large pots: 38 cm length × 25 cm width) (‘CPP-Brötchenbeutel genadelt’, www.der-verpackungs-profi.de GmbH, Göttingen, Germany) was placed over the plant and was secured with a rubber band around the pot. Inoculated plants in small and large pots were kept in two different environments: (A) 14:10 h light/dark photoperiod, 24:18 °C, 60-80% r.h., 200 µmol (s m²)^-1^ (‘Valoya 03-155-230 R150 AP673 LED Oberlicht’, Valoya Oy, Helsinki, Finland) and (B) 16:8 h light/dark photoperiod, 20.9 ± 1 °C, 48 ± 12.2% r.h. and 80 µmol (s m²)^-1^ of full-spectrum light (described above). Egg batches were removed and quantified 16-18 days after inoculation. Each treatment was tested with six replicates.

### 2.4 Role of host plant species for adult survival and oviposition

The role of sugar beet and wheat plants for survival and oviposition in *P. leporinus* was assessed. Sugar beet (BBCH 12-14, n = 17) and two growth stages of wheat plants *Triticum aestivum* cv. Dekan (KWS SAAT SE & Co. KGaA, Einbeck, Germany) were compared in a no-choice experiment (‘small wheat plants’: BBCH 10-14, n = 15 and ‘large wheat plants’: BBCH 19, n = 9).^21^ Plants were cultivated in small pots with a single sugar beet or 2-3 wheat plants per pot. Sugar beet and small wheat plants were grown under controlled climatic conditions (see 2.2). Large wheat plants were grown in a greenhouse before the experiment (see 2.1). All plants were supplied with a single, newly emerged (max. 24 h old) female and 1-2 male adult planthoppers and covered with a transparent, perforated polypropylene bag (25 cm length × 15 cm width). Plants were arranged in a complete randomized design in the controlled environment room. Oviposition of adults was evaluated daily and survival of adults was evaluated every third day. Egg batches were removed from the substrate and the number of eggs was counted under a stereomicroscope (model ‘M3Z’, Wild Heerbrugg AG, Heerbrugg, Switzerland).

### 2.5 Continuous rearing of *Pentastiridius leporinus*

For oviposition, five female and three male adult planthoppers were kept on a single potted sugar beet plant (BBCH 12-14).^21^ Plants were grown in small pots and covered with a perforated bag (Fig. 4A). Over a 4-6 week period, egg batches were removed after seven days (Fig.4B) and groups of 4-5 egg batches were transferred to a plastic container (18 cm length × 13.5 cm width × 6 cm height) (Salatschale NP eckig - Becher - Polystyrol weiß - 1000 g, Papier Brinkmann GmbH, Münster, Germany), which contained a 1 cm layer of 3:1 peat-sand mix (Fig. 4C) and was kept in darkness by placing eight containers in two inverted plastic trays (60 cm length × 40 cm width × 7.5 cm height) (Newbox 15, beku Lagertechnik GmbH, Klagenfurt, Austria). The container was closed with a transparent lid that contained holes for gas exchange. The substrate surrounding the egg batches was kept moist by adding a few drops of water once per week until nymphs hatched to prevent eggs from desiccating, whereas the rest of the substrate inside the box was kept dry. This measure was taken as several nymphs had died and expressed mycosis when humidity in the container was too high. Bisected or quartered taproots from sugar beet (BBCH 19: 10 to 20 unfolded leaves), grown in polypropylene pots (9 cm length × 9 cm width × 9.5 cm height) in a greenhouse, were added as a food source for the hatching nymphs (Fig. 4D). Taproots were replaced after 14 days or earlier when showing signs of deterioration. Every second week, rearing containers were checked for fifth instar nymphs (first appearance after ca. eight weeks). All fifth instars were gently removed with a laboratory spoon (Fig. 4E), transferred to another container and released into an insect rearing tent with potted sugar beet plants (large pots with a single plant in BBCH 16-19: 6 to 14 unfolded leaves) to complete metamorphosis on an intact plant (Fig. 4F). Large pots were inoculated with 100 fifth instar nymphs. The substrate consisted of two layers with a 3:1 peat-sand mix as the lower part and a top layer of approx. 5 cm expanded clay (‘Original LamstedtTon 8–16 mm’, Fibo ExClay Deutschland GmbH). Cages were checked once per week for emerged adult planthoppers. Adults were gently removed with an aspirator (Fig. 4G), sexed and used for oviposition as described above or for further experiments.

### 2.6 Characterisation of developmental stages

In Central Europe, winter wheat is the most common crop grown after sugar beet harvest. Accordingly, *P. leporinus* nymphs spend most of their development on wheat roots until reaching the imago stage.^4^ To measure the developmental time of immature planthoppers, a single nymph was placed in a plastic container (11 cm length × 8 cm width × 5 cm height) (‘Saladboxx 250 cc’, Pro-Pac Ostendorf Plastic Thermoformfolien und Verpackungen GmbH & Co. KG, Vechta, Germany) with small holes in the lid for gas exchange. The container was filled with a layer of 3:1 peat-sand mix (25 g), which was kept moist and replaced after ca. 4 weeks. Two 4-10 days old wheat seedlings were added as a food source and replaced after eight days or earlier, if needed. The entire experiment was carried out twice. In the first replicate, the offspring of at least seven different parental females from ten different egg batches were analysed (4-14 nymphs per egg batch) and recordings started with second instar nymphs (n = 78) and ended with the fifth instar. Until reaching the second stage, neonates were kept on sugar beet. In the second replicate, the entire life cycle for the offspring of 12 different parental females (8-12 nymphs per female) was assessed. This experiment commenced by taking egg batches with known oviposition dates and transferring these to Petri dishes. At the day of hatching, first instar nymphs (n = 105) were transferred to the described experimental setup using a small paint brush. Survival and development of the nymphs were recorded every day under a stereomicroscope.

Morphometric measurements of each nymphal instar and of eggs from three different generations were taken to determine if body size can be used to differentiate between juvenile stages. Maximum length and width of eggs was recorded with a stereomicroscope (40x magnification). Body length of nymphal instars was measured (10-40x magnification) from the tip of the vertex to the tip of the abdomen.^22^ Head capsule width was determined as indicated in Fig. 1.

**Figure 1.**
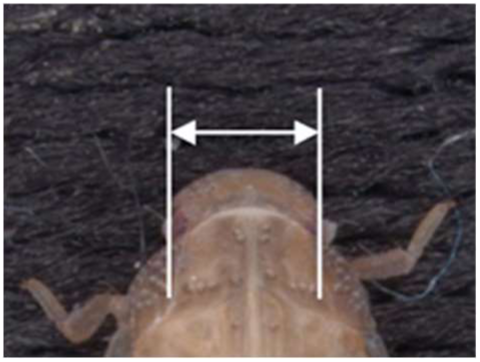
Anterior part of *Pentastiridius leporinus* nymph. Lines and arrow indicate how head capsule width was determined.

Photos of the different developmental stages were taken with focus stacking under a stereomicroscope model ‘MSV266’ to which a camera model ‘DMC5400’ (both Leica Microsystems GmbH, Wetzlar, Germany) was fitted.

### 2.7 Infection rate of planthoppers

To examine the infection rate with SBR pathogens in the rearing, randomly selected adult *P. leporinus* (n = 10) from three consecutive generations were analysed with nested polymerase chain reaction (PCR). Insects were individually stored in 96% (v/v) ethanol at -20 °C until use. DNA was extracted with DNeasy® Blood & Tissue Kit (QIAGEN GmbH, Hilden, Germany) according to the manufacturer’s instructions. Presence of *Ca*. Arsenophonus phytopathogenicus was verified with the primers Fra5/L1r and Alb1/Oliv1 as described in Semetey et al. (2007), infection with the stolbur phytoplasma was tested with the primers P1/P7 and U5/U3 as described in Gatineau et al. (2001).^16,23^

### 2.8 Statistical analyses

All statistical analyses were carried out with SAS 9.4 (SAS Institute Inc., Cary, NC, USA). In the oviposition experiment, data were analysed separately for both environments. Two-way analysis of variance (ANOVA) was carried out after testing for homogeneity of variances and normal distribution. Accordingly, oviposition data in the environment (b) were square-root transformed to meet the condition of variance homogeneity. Least square means were calculated to analyse significant differences at p<0.05 using ‘lsmeans’ in the PROC GLIMMIX procedure.^24^

Survival analysis of the no-choice oviposition experiment was carried out with Cox proportional hazard models and PROC PHREG in SAS 9.4. Hazard ratios (HR) >1 or <1 describe higher or lower probabilities of mortality compared to the referred treatment, respectively.^24^

## 3 RESULTS

### 3.1 Role of plant container size, substrate and environmental conditions for oviposition success

Egg batches were produced in all provided plant containers, substrates, and environments but significant differences were found among treatments (Fig. 2). In both environments, the highest number of egg batches (≥ 1.5) per female was found in large pots with field soil. Significantly fewer egg batches were found in pots with other substrates, although no clear pattern could be discerned with regard to the role of pot size and substrate. In environment A, only small pots with field soil contained a median of more than 1 egg batch per female (Fig. 2A).

**Figure 2.**
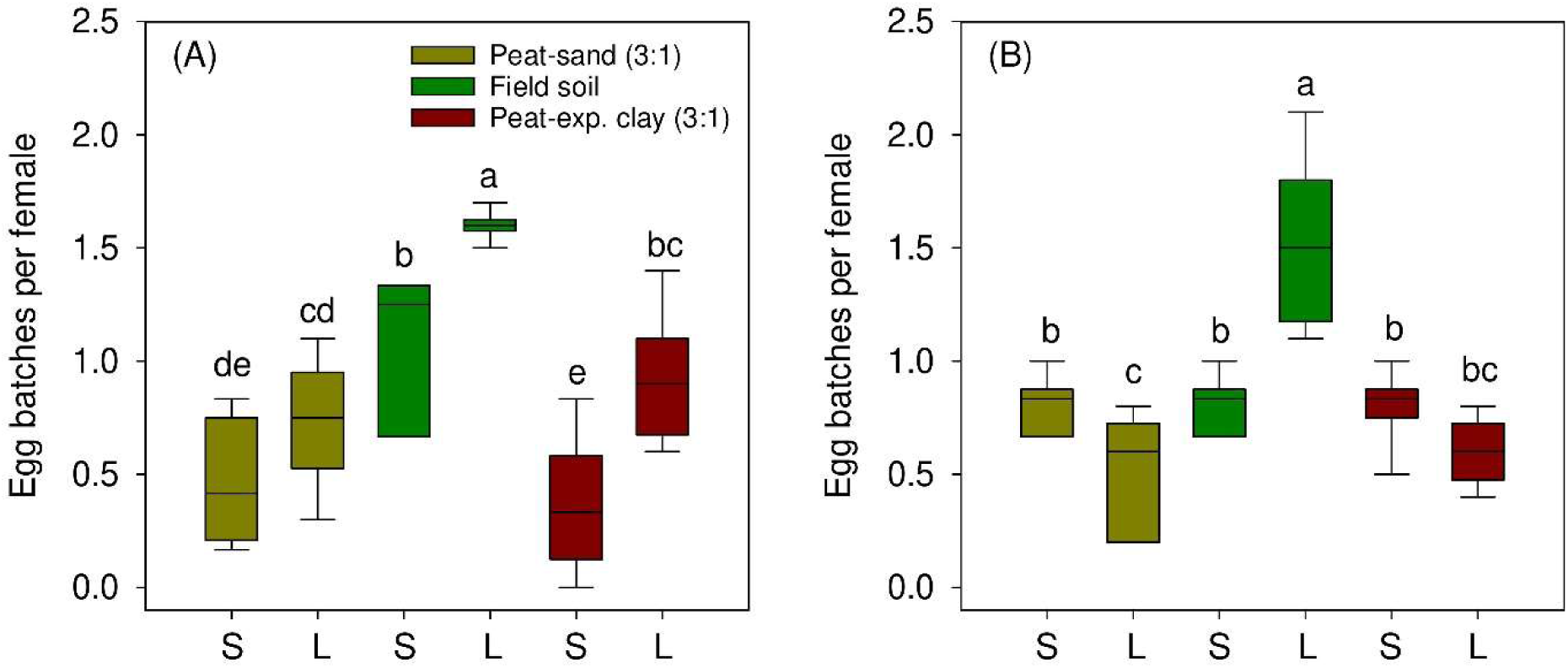
Oviposition of *Pentastiridius leporinus* in potted sugar beet in dependence of pot type, substrate and two environments: (A) 14:10 h light/dark photoperiod, 24:18 °C, 60-80% r.h. and 200 µmol (s m²)^-1^ and (B) 16:8 h light/dark photoperiod, 20.9 ± 1 °C, 48 ± 12.2% r.h. and 80 µmol (s m²)^-1^. Small pots (S) with 4.8 cm diameter and large pots (L) with 16 cm diameter were compared (n = 6). Boxes show 25^th^, 50^th^ and 75^th^ percentiles, whiskers show 10^th^ and 90^th^ percentiles. Treatments with the same letter within a graph are not significantly different according to two-way ANOVA (α = 0.05).

No significant differences were found between large pots filled with either peat-sand or peat-expanded clay substrate. The same was true for small pots with both peat substrates. In environment B, no differences in the number of egg batches (median 0.81) were detected between different substrates in small pots (Fig. 2B). Interestingly, large pots with either peat substrate contained fewer or similar numbers of egg batches than small containers.

### 3.2 Role of host plant species for adult survival and oviposition

The mortality rates of female adults kept on small (HR: 13.6, Cox proportional hazard model, p < 0.001) and large (HR: 13.2, Cox proportional hazard model, p < 0.001) wheat plants were significantly higher compared to female adults kept on sugar beet (Fig. 3). No statistical differences in mortality were observed on small compared to large wheat plants (HR: 1.04, Cox proportional hazard model, p = 0.93). After 18 days, all planthoppers had died in the wheat treatments, while insects in the sugar beet treatment survived for 53 days.

**Figure 3.**
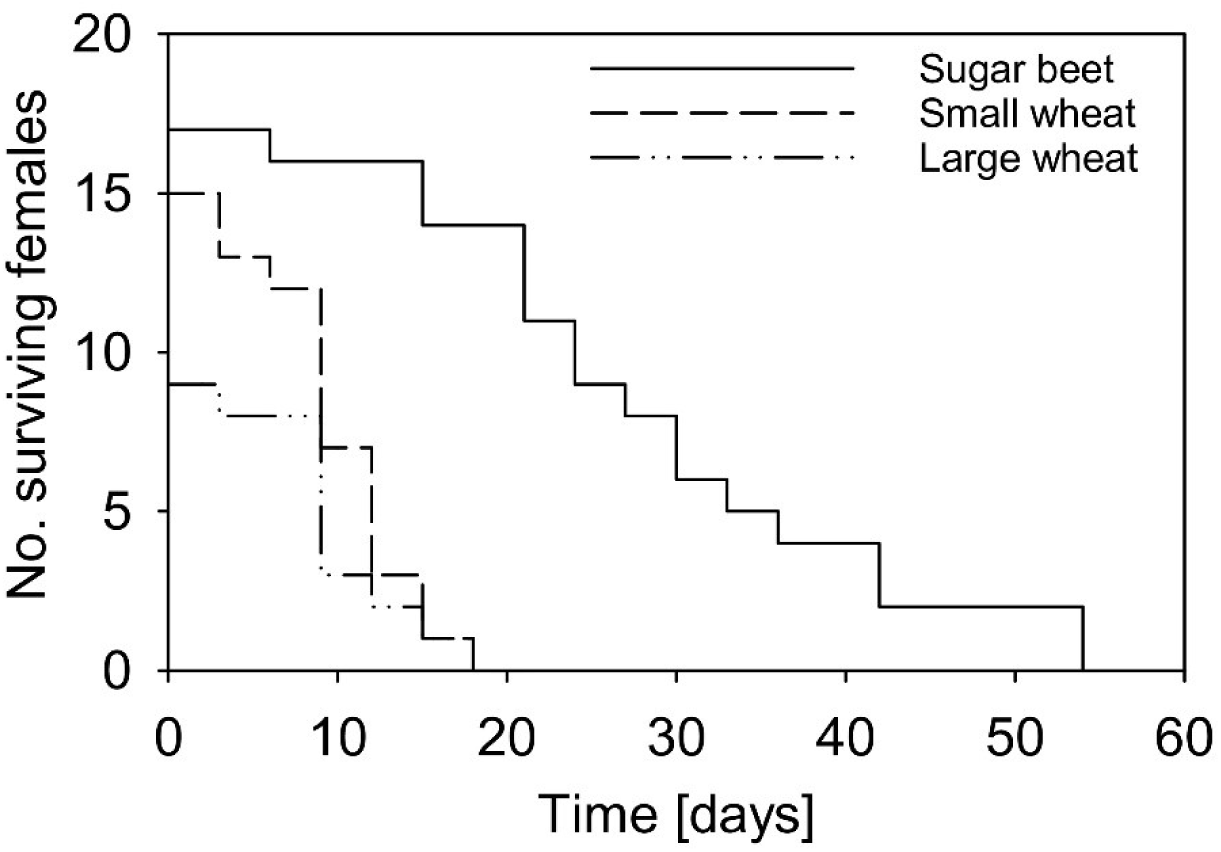
Survival of single female adult *Pentastiridius leporinus* on sugar beet (n = 17), small (BBCH 10-14, n = 15) and large wheat plants (BBCH 19, n = 9) in a no-choice experiment.

Neither the production of wax filaments on the tip of the female’s abdomen, a characteristic trait of ovipositing *P. leporinus*, nor any egg batches were found in the wheat treatments. In contrast, a maximum of eight oviposition events per planthopper was observed in sugar beet (Table 1).

**Table 1.** Temporal oviposition pattern of *Pentastiridius leporinus* in a no-choice experiment. Data show time after eclosion when egg batches were laid and number of eggs per egg batch for each oviposition event of a single female. Oviposition occurred only in sugar beet plants (n = 17), no eggs were laid in wheat.

| Oviposition event | Days |  |  | Eggs per batch |  |  | n |
| --- | --- | --- | --- | --- | --- | --- | --- |
| | Range | Median | Mean±SD | Range | Median | Mean±SD | |
| 1 | 6-17 | 11 | 10.6±3.5 | 33-186 | 51 | 51±44.9 | 13 |
| 2 | 10-25 | 17 | 17±4 | 0-77 | 42 | 42±21.2 | 11 |
| 3 | 11-24 | 19 | 18.8±4.8 | 0-48 | 40 | 40±19.8 | 6 |
| 4 | 14-32 | 29 | 25.8±7.2 | 18-77 | 37 | 37±21.7 | 6 |
| 5 | 18-36 | 27.5 | 28.3±7.5 | 10-66 | 17 | 17±30.5 | 4 |
| 6 | 33-42 | 37.5 | 37.5±6.4 | 8-38 | 23 | 23±21.2 | 2 |
| 7 | - | 33 | 33 | - | 42 | 42 | 1 |
| 8 | - | 39 | 39 | - | 39 | 39 | 1 |

In total, 13 of 17 female adults laid eggs and more than six oviposition events were observed in a single female. Females were observed to lay eggs up to 42 days after eclosion. The mean number of eggs per egg batch was 46.9 ± 32.5 with most eggs per egg batch being laid in the first oviposition event. The largest batch consisted of 186 eggs. A mean of 139.1 ± 132.9 and a maximum of 406 eggs per female adult were observed.

### 3.3 Continuous rearing of *Pentastiridius leporinus*

A continuous mass rearing of *P. leporinus* was successfully established. The rearing started with field collected planthoppers in June 2020 and until March 2022 five generations and >20,000 *P. leporinus* individuals were produced under controlled climatic conditions. Crucial points were the transfer of fifth instar nymphs to potted sugar beets with a top layer of expanded clay for metamorphosis (Fig. 4F) to avoid the development of adults inside the containers (Fig. 4C-E). This step prevented losing adult planthoppers due to starvation as they are probably unable to feed on roots. The nymphs immediately hid between the expanded clay particles after being transferred. Without a top layer of expanded clay metamorphosis failed, as most transferred nymphs left the pot and died within a few days (data not shown). Adult emergence started approximately 4 weeks after the transfer (Fig. 5) and 140 days later 68.7% of the nymphs had developed into adults. The majority of emerging adults were male (59.2%). After 140 days, the recording was stopped since adult emergence had nearly ceased.

**Figure 4.**
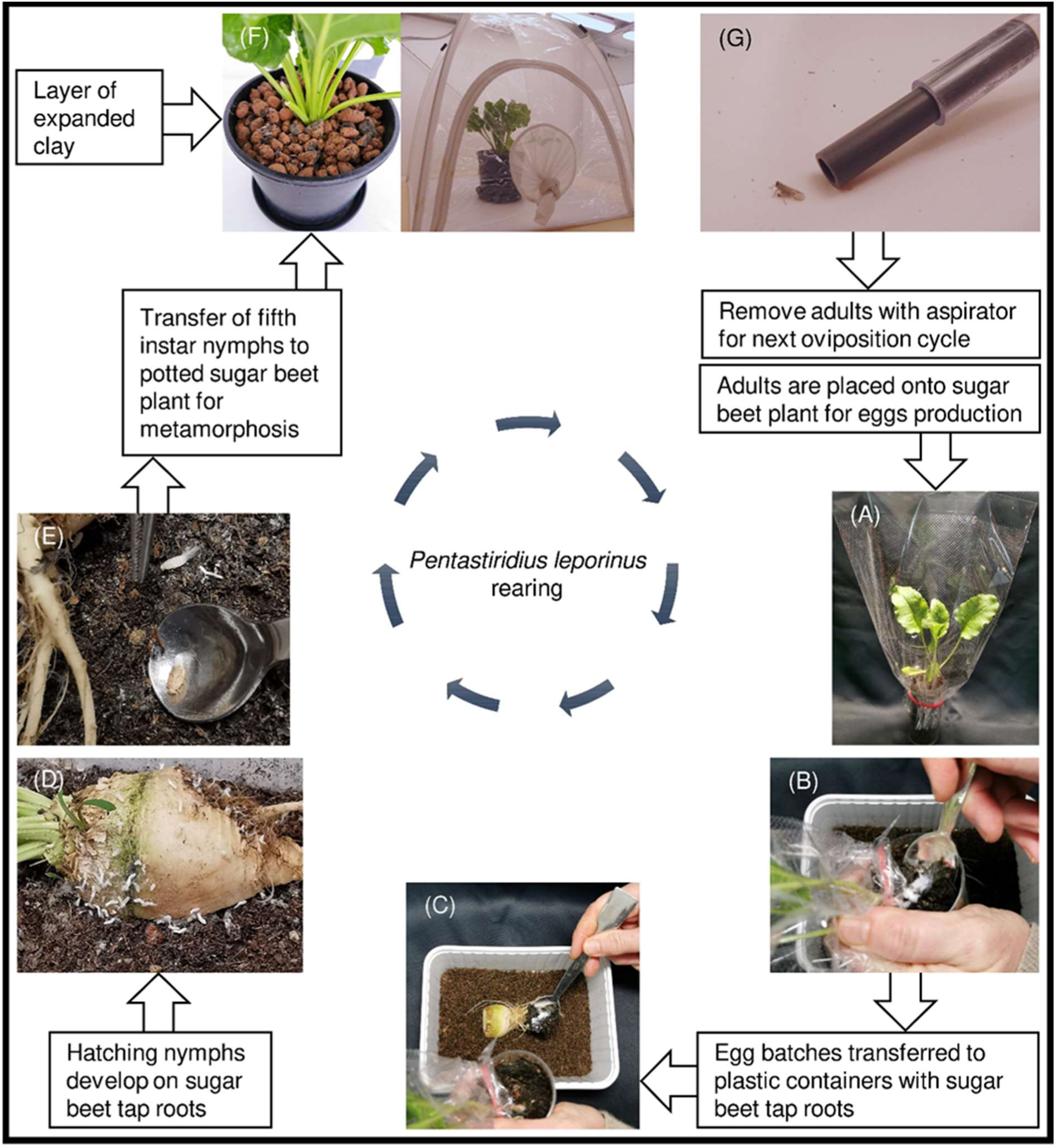
Schematic representation of laboratory method for *Pentastiridius leporinus* rearing.

### 3.4 Characterisation of developmental stages

The highest mortality (25.7%) was observed in first instar nymphs (Table 2). Lowest mortality (0-5.9%), on the other hand, was found in third and fourth nymphal instars. In total, 56.2% of the analysed specimens from the second generation completed their life cycle on wheat seedlings. With older instars, the average and range of developmental time increased. First female and male adults emerged 104 and 145 days after hatching. Body length and head capsule width increased with larger nymphal instars (Table 3). In contrast to body length, head capsule width did not overlap between nymphal instars. All nymphal instars produced wax filaments. All developmental stages are depicted in Fig. 6.

**Table 2.** Larval mortality and duration of developmental stages of *Pentastiridius leporinus* reared on wheat plants at 20.9 ± 1 °C. Insects from two generations were analysed. n = number of individuals with known beginning and ending of a developmental stage. SD = Standard deviation. Range indicates minimum and maximum numbers of days per stage.

| Stage | Mortality [%] | Days |  |  | n |
| --- | --- | --- | --- | --- | --- |
|  |  | Range | Median | Mean±SD |  |
| Egg batch | - | 17-25 | 22 | 21.6±1.5 | 52 |
| First instar | 25.7 | 13-21 | 16 | 16.9±2.1 | 39 |
| Second instar | 12.9 | 13-27 | 16 | 16.9±3.5 | 29 |
| Third instar | 3.0 | 15-48 | 22 | 24.6±8.3 | 55 |
| Fourth instar | 3.1 | 19-151 | 30 | 48.9±36.2 | 45 |
| Fifth instar | 13.4 | 19-201 | 97 | 105.7±48.0 | 66 |
| First instar-male adult | - | 145-264 | 185 | 193.6±35.8 | 31 |
| First instar-female adult | - | 104-354 | 181 | 193.5±59.2 | 24 |

**Table 3.** Morphometric measurements showing length and width of egg and nymphal stages of *Pentastiridius leporinus*. In nymphs, width was measured from head capsules.

| Stage | Length [mm] |  | Width [mm] |  | n |
| --- | --- | --- | --- | --- | --- |
|  | Range | Mean ± SD | Range | Mean ± SD |  |
| Egg | 0.57-0.74 | 0.65±0.05 | 0.28-0.36 | 0.33±0.02 | 25 |
| First instar | 0.74-1.24 | 0.92±0.13 | 0.26-0.29 | 0.27±0.01 | 30 |
| Second instar | 1.11-1.76 | 1.43±0.18 | 0.31-0.40 | 0.35±0.02 | 32 |
| Third instar | 1.60-2.41 | 2.03±0.22 | 0.43-0.57 | 0.48±0.03 | 52 |
| Fourth instar | 1.80-4.20 | 2.83±0.48 | 0.60-0.78 | 0.69±0.04 | 102 |
| Fifth instar | 2.76-5.93 | 4.26±0.67 | 0.85-1.22 | 0.98±0.08 | 69 |

**Figure 5.**
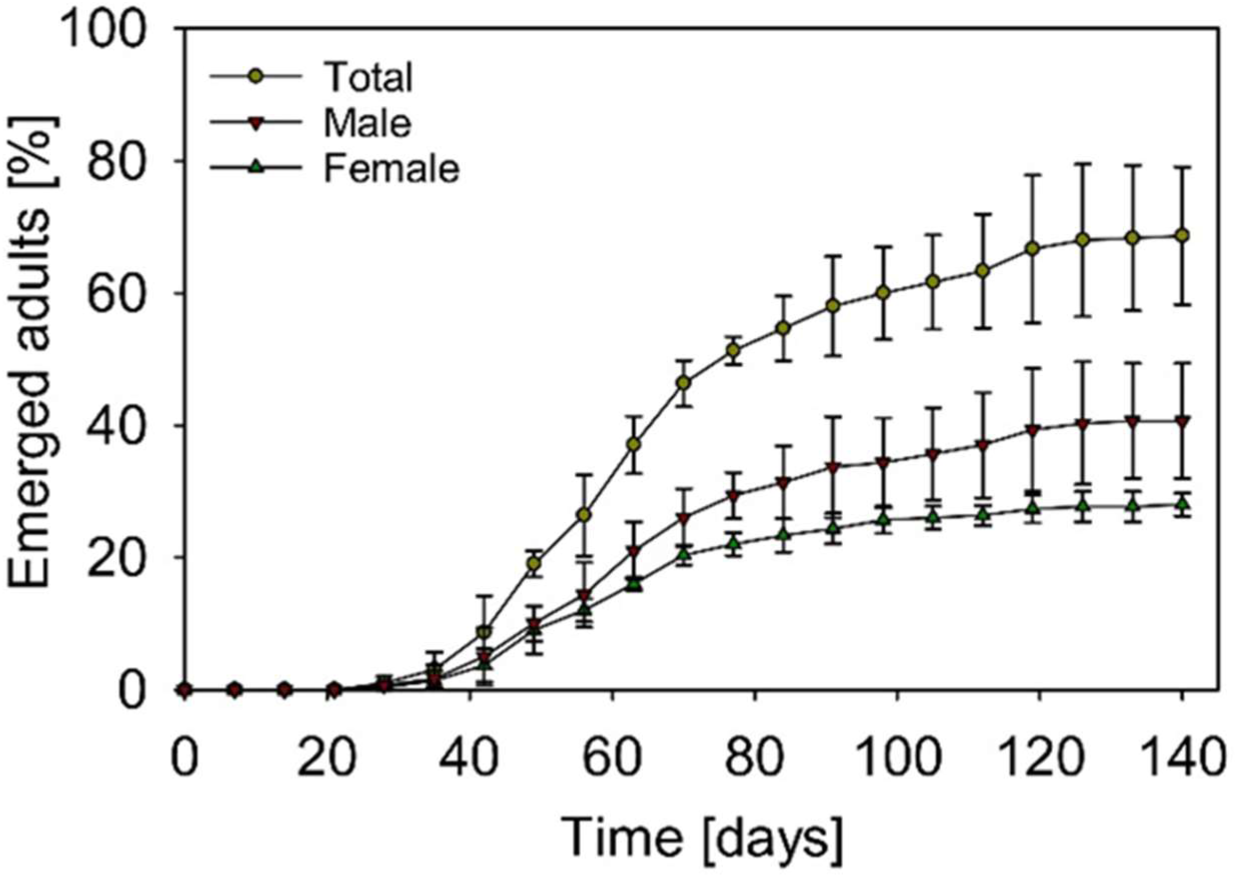
Accumulated percentage of emerged adults from fifth instar nymphs on potted sugar beet plants which were grown in large pots with a top layer of expanded clay (n = 3). Symbols show mean values, whiskers show standard deviation.

**Figure 6.**
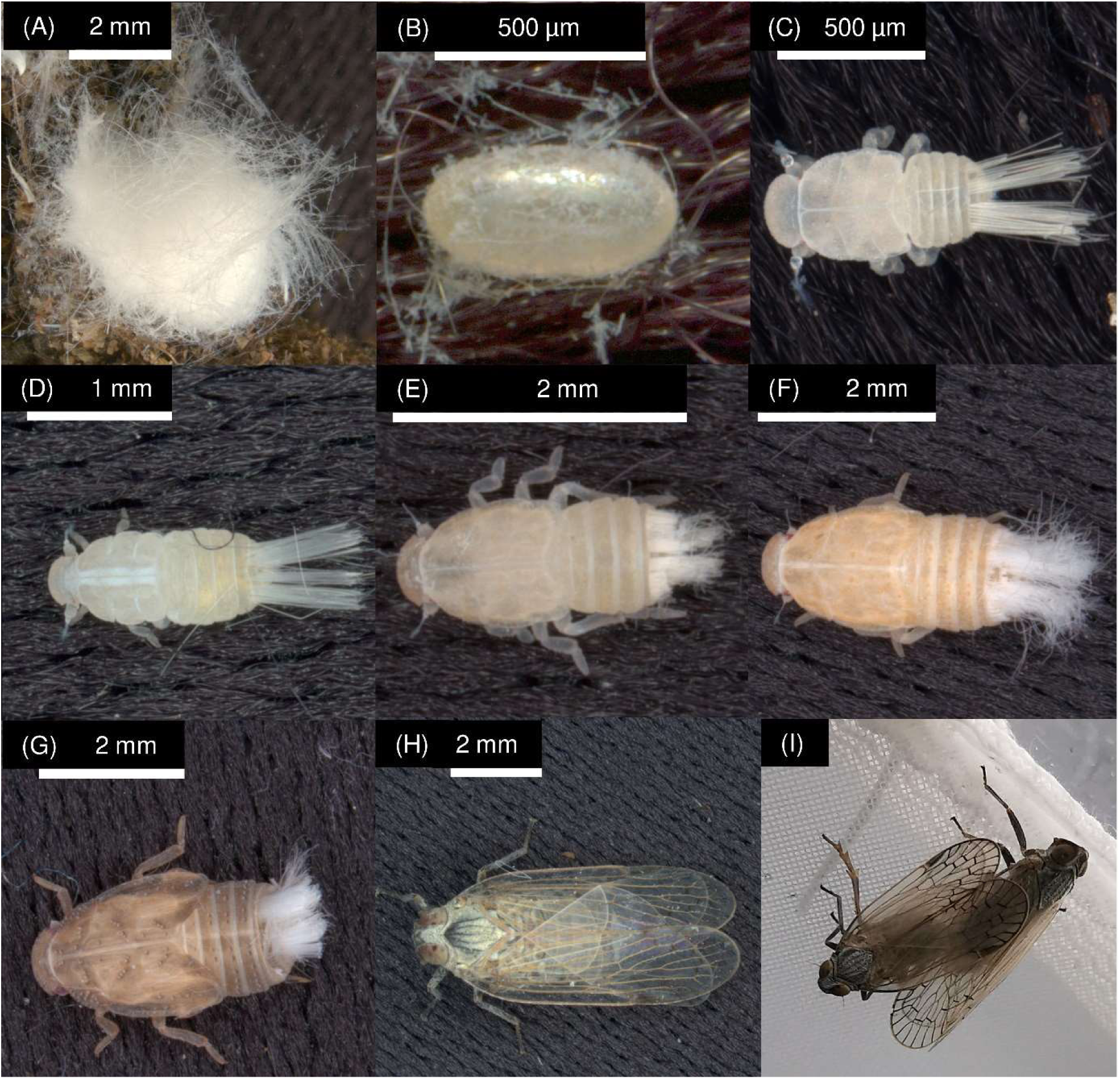
Development stages of *Pentastiridius leporinus*. **(A)** Egg batch covered with waxy filaments, **(B)** single egg, **(C)** first instar nymph, **(D)** second instar nymph, **(E)** third instar nymph, **(F)** fourth instar nymph, **(G)** fifth instar nymph, **(H)** female adult and **(I)** copulation.

### 3.5 Infection rate of adult planthoppers

Adults infected with the proteobacterium were detected in all three generations of the rearing (Table 4). The infection rates (70-80%) were comparable between the different generations. In contrast, no phytoplasma was detected in any of the analysed adults.

**Table 4.** Infection rate of adult *Pentastiridius leporinus* with the γ-3 proteobacterium (Proteo) *Ca*. Arsenophonus phytopathogenicus and phytoplasma (Phyto) *Ca*. Phytoplasma solani in different generations of the rearing. The ratio of positive to total tested specimens is shown.

|  | 1<sup>st</sup> generation |  | 2<sup>nd</sup> generation |  | 3<sup>rd</sup> generation |  |
| --- | --- | --- | --- | --- | --- | --- |
|  | Proteo | Phyto | Proteo | Phyto | Proteo | Phyto |
| Infection rate | 7/10 | 0/10 | 8/10 | 0/10 | 8/10 | 0/10 |

## 4 DISCUSSION

Here we describe a new protocol for the continuous mass rearing of the planthopper *P. leporinus*. Our study also provides basic life history data on this little-known insect species and a simple method for distinguishing nymphal stages.

As a first step towards developing a suitable rearing method and to obtain an idea which plant container size, substrate or combination of environmental conditions would yield a high number of egg batches per female, an oviposition experiment was performed with field-collected adult planthoppers. We found that it was possible to produce *P. leporinus* eggs in two different environmental settings that represented climate chamber conditions common for rearing insects of the temperate zone. The combination of small pots with six female planthoppers versus large pots with ten female planthoppers did not yield a clear picture, with one exception: the highest number of egg batches was always found in large pots containing field soil. In these pots, egg batches were mostly detected in the gap between substrate and pot wall. Field soil contained many cracks and cavities, while both alternative substrates were rather compacted. Field soil therefore allowed *P. leporinus* females to walk into the cracks for oviposition, which is relevant as *P. leporinus* has not been observed to dig. Although large pots filled with field soil turned out to be superior compared to all other combinations, the important information gained in this trial was that *P. leporinus* also accepted small transparent polypropylene pots with standardized, commercially available substrates for oviposition. The small size of the containers allowed easy handling, while the transparent polystyrene made it possible to regularly monitor the presence of egg batches. Hence, these options were chosen for further mass rearing.

In a second trial, we studied the oviposition pattern and longevity of *P. leporinus* on sugar beet and wheat plants. The results were unequivocal as no eggs were laid at all in pots with wheat and planthoppers feeding on this host plant lived only for around 18 days. Also, no abdominal wax filaments were produced by females caged onto wheat. Such wax filaments were generally found in mature females about to lay eggs in the sugar beet treatment and are a characteristic feature in several cixiid planthoppers.^1,25,26^ Females offered sugar beet, on the other hand, produced several egg batches with an average of 139 eggs during their entire life span of 53 days. These egg batches contained a mean of 47 eggs. The results are interesting because wheat is a very suitable host for nymphal development and hence the question of why *P. leporinus* females reject this species as an oviposition substrate awaits further exploration. A falsifiable hypothesis could be that wheat phloem sap cannot be exploited by adult *P. leporinus* as an appropriate food source that supports egg production.

In the sugar beet – winter wheat system, nymphs spend most of their life on wheat roots.^4^ Hence, the developmental time and mortality of nymphal stages was recorded on this host. We found that highest mortality occurred in first instars at a rate of 25.7% and declined strongly in instars three and four. However, fifth instar nymphs were more vulnerable again, accounting for 10.6% in mortality. Remarkably, not only developmental time increased consistently with each nymphal instar but also the time range that nymphs spent in the respective stages. While the first instar stage lasted 13-21 days, fifth instars varied considerably between 17 and 211 days. Overall, male and female planthoppers had the same lifespan and the whole life cycle was completed in approximately seven months under laboratory conditions. Additionally, morphometric measures of all immature stages were taken. Head capsule width was thereby identified as a trait that lends itself well to unequivocally differentiate between nymphal instars, as no overlap was found for head capsule widths.

With the gained insights, we were able to establish a continuous mass rearing as described above and shown in Fig. 4. We exclusively used sugar beet as a host, which furthermore demonstrates that *P. leporinus* can complete its life cycle entirely on this plant without the need for alternative hosts such as wheat. From June 2020 until March 2022, we were able to produce five generations and over 20,000 nymphs under controlled climatic conditions at constant 20.9 ± 1°C. Obviously, *P. leporinus* does not require an obligatory diapause, although the species is known to have a univoltine life cycle in Central Europe.^2,3^ As a crucial step in our rearing, we found that it was necessary to provide fifth instars with potted sugar beet plants that had a 5 cm layer of expanded clay on top of the peat-sand substrate. This allowed the insects to gain access to the roots and complete their metamorphosis. Attempts to produce adults on sugar beet plants without this top layer of expanded clay had failed. In our experiments, 68.7% of fifth instar nymphs emerged as adults within 140 days of which the majority (59.2%) were male. So far, we are unaware of any other protocol for mass rearing of Cixiidae with belowground nymphal stages. Sforza et al. (1999) reported the rearing of epigean nymphs of the related species *H. obsoletus*. However, mass rearing was not possible due to high mortality of the insects.^22^

Adult planthoppers from three consecutive generations were taken from the rearing and tested for the presence of *Ca*. Arsenophonus phytopathogenicus and *Ca*. Phytoplasma solani by nested PCR. While the sample size was comparatively low, the emerging picture was clear, showing that 70-80% of insects from three generations were infected with the proteobacterium. The phytoplasma was absent in all 30 specimens tested. Bressan et al. showed that vertical transmission of the SBR proteobacterium to the next generation is possible.^18^ This finding could be confirmed in our studies because the three generations of *P. leporinus* were held separately. In summary, our study provides relevant information on several life history traits of *P. leporinus* and introduces a method for its continuous mass rearing. The presented data will facilitate research on the biology and ecology of this economically important vector insect and can thus critically contribute to the development of integrated pest management strategies.

## Acknowledgements

We thank Bianca Tappe, Chelsea Schreiber, Gülsen Aydin, Jonas Watterott, Jutta Schaper, and Katrin Waldstein (Division of Agricultural Entomology, Göttingen) as well as Annette Tostmann, Annette Walter, Celin Lachmann, and Georgia Hesse (Institute of Sugar Beet Research, Göttingen) for technical assistance. This research was funded by Kuratorium für Versuchswesen und Beratung im Zuckerrübenanbau (Mannheim, Germany), the Institute of Sugar Beet Research and the Division of Agricultural Entomology.

## Data availability statement

Research data are not shared.

## References

1 Holzinger WE, Emeljanov AF and Kammerlander I, Zikaden - leafhoppers, planthoppers and cicadas. The family Cixiidae Spinola 1839 (Hemiptera, Fulgoromorpha) – a review. Denisia 4:113–138 (2002).

2 Holzinger WE, Kammerlander I and Nickel H, The Auchenorrhyncha of Central Europe - Die Zikaden Mitteleuropas: Volume 1: Fulgoromorpha, Cicadomorpha excl. Cicadellidae, Brill NV, Leiden, Netherlands (2003).

3 Biedermann R and Niedringhaus R, Die Zikaden Deutschlands: Bestimmungstafeln für alle Arten, Wissenschaftlicher Akademischer Buchvertrieb, Fründ, Scheeßel, Germany (2004).

4 Bressan A, Moral García FJ, and Boudon-Padieu E, The prevalence of ‘Candidatus Arsenophonus phytopathogenicus’ infecting the planthopper Pentastiridius leporinus (Hemiptera: Cixiidae) increase nonlinearly with the population abundance in sugar beet fields. Environ Entomol 40:1345–1352 (2011). https://doi.org/10.1603/EN10257.

5 Bressan A, Terlizzi F and Credi R, Independent origins of vectored plant pathogenic bacteria from arthropod-associated Arsenophonus endosymbionts. Microb Ecol 63:628–638 (2012). https://doi.org/10.1007/s00248-011-9933-5.

6 Gatineau F, Jacob N, Vautrin S, Larrue J, Lherminier J, Richard-Molard M and Boudon-Padieu E, Association with the syndrome “basses richesses” of sugar beet of a phytoplasma and a bacterium-like organism transmitted by a Pentastiridius sp. Phytopathology 92:384–392 (2002). https://doi.org/10.1094/PHYTO.2002.92.4.384.

7 Peter M, Neue Krankheit bedroht den Zuckerrübenanbau, LANDfreund 1:24–25 (2020).

8 Pfitzer R, Schrameyer K, Voegele RT, Maier J, Lang C and Varrelmann M, Causes and effects of the occurrence of “Syndrome des basses richesses” in German sugar beet growing areas, Sugar Industry 145:234–244 (2020). https://doi.org/10.36961/si24263

9 Sémétey O, Bressan A, Richard-Molard M and Boudon-Padieu E, Monitoring of proteobacteria and phytoplasma in sugar beets naturally or experimentally affected by the disease syndrome ‘Basses richesses’, Eur J Plant Pathol, 117:187–196 (2007). https://doi.org/10.1007/s10658-006-9087-3.

10 Bressan A, Agronomic practices as potential sustainable options for the management of Pentastiridius leporinus (Hemiptera: Cixiidae) in sugar beet crops. J Appl Entomol 133:760–766 (2009). https://doi.org/10.1111/j.1439-0418.2009.01407.x.

11 Bressan A, Holzinger WE, Nusillard B, Sémétey O, Simonato M and Boudon-Padieu E, Identification and biological traits of a planthopper from the genus Pentastiridius (Hemiptera: Cixiidae) adapted to an annual cropping rotation. Eur J Entomol 106:405–413 (2009). https://doi.org/10.14411/eje.2009.052.

12 Nickel H, Achtziger R, Biedermann R, Bückle C, Deutschmann U, Niedringhaus R, Remane R, Walter S and Witsack W, Rote Liste und Gesamtartenliste der Zikaden (Hemiptera: Auchenorrhyncha) Deutschlands, in: Rote Liste gefährdeter Tiere, Pflanzen und Pilze Deutschlands, Band 4: Wirbellose Tiere (Teil 2), ed. by Gruttke H, Balzer S, Binot-Hafke M, Haupt H, Hofbauer N, Ludwig G, Matzke-Hajek G and Ries M Münster (Landwirtschaftsverlag), Naturschutz und Biologische Vielfalt 70:249–298 (2016).

13 Bressan A, Sémétey O, Nusillard B, Clair D and Boudon-Padieu E, Insect vectors (Hemiptera: Cixiidae) and pathogens associated with the disease syndrome “basses richesses” of sugar beet in France. Plant Dis 92:113–119 (2008). https://doi.org/10.1094/PDIS-92-1-0113.

14 Schröder M, Rissler D and Schrameyer K, „Syndrome des Basses Richesses” (SBR) – erstmaliges Auftreten an Zuckerrüben in Deutschland, J Cultivated Plants 64:396–397 (2012).

15 Firrao G, Gibb K and Streten C, Short taxonomic guide to the genus ‘Candidatus Phytoplasma’. J Plant Pathol, 87:249–263 (2005).

16 Gatineau F, Larrue J, Clair D, Lorton F, Richard-Molard M and Boudon-Padieu E, A new natural planthopper vector of stolbur phytoplasma in the genus Pentastiridius (Hemiptera: Cixiidae). Eur J Plant Pathol, 107:263–271 (2001). https://doi.org/10.1023/A:1011209229335.

17 Sémétey O, Gatineau F, Bressan A and Boudon-Padieu E, Characterization of a γ-3 proteobacteria responsible for the syndrome “basses richesses” of sugar beet transmitted by Pentastiridius sp. (Hemiptera, Cixiidae), Phytopathology 97:72–78 (2007). https://doi.org/10.1094/PHYTO-97-0072.

18 Bressan A, Sémétey O, Arneodo J, Lherminier J and Boudon-Padieu E, Vector transmission of a plant-pathogenic bacterium in the Arsenophonus clade sharing ecological traits with facultative insect endosymbionts. Phytopathology 99:1289–1296 (2009). https://doi.org/10.1094/PHYTO-99-11-1289.

19 Bressan A, Emergence and evolution of Arsenophonus bacteria as insect-vectored plant pathogens. Infect, Genet Evol 22:81–90 (2014). https://doi.org/10.1016/j.meegid.2014.01.004.

20 Mühlenberg M, Freilandökologie, Quelle & Meyer, Heidelberg/Wiesbaden, Germany (1993).

21 Meier U, Growth Stages of Mono-and Dicotyledonous Plants. BBCH Monograph, Federal Biological Research Centre for Agriculture and Forestry, Berlin/Braunschweig, Germany (2001).

22 Sforza R, Bourgoin T, Wilson SW and Boudon-Padieu E, Field observations, laboratory rearing and descriptions of immatures of the planthopper Hyalesthes obsoletus (Hemiptera: Cixiidae), Eur J Entomol 96:409–418 (1999). https://www.eje.cz/pdfs/eje/1999/04/15.pdf

23 Sémétey O, Bressan A, Gatineau F and Boudon-Padieu E, Development of a specific assay using RISA for detection of the bacterial agent of ‘basses richesses’ syndrome of sugar beet and confirmation of a Pentastiridius sp. (Fulgoromopha, Cixiidae) as the economic vector, Plant Pathol 56:797–804 (2007). https://doi.org/10.1111/j.1365-3059.2007.01693.x.

24 SAS Institute Inc., SAS/STAT® User’s Guide. SAS Institute Inc., Cary, NC, USA (2022).

25 Cumber RA, Studies on Oliarus atkinsoni Myers (Hem. Cixiidae), vector of the “yellow leaf” disease OF Phormium tenax Forst. NZ J Sci Technol 34:92–98 (1952).

26 Mason RT, Fales HM, Jones TH, O’Brien LB, Taylor TW, Hogue CL, Blum MS, Characterization of fulgorid waxes (Homoptera:Fulgoridae:Insecta). Insect Biochem 19:737–740 (1989). https://doi.org/10.1016/0020-1790(89)90054-1.

